# Metabolic allometry: A genomic approach to scaling

**DOI:** 10.1101/2020.12.27.424503

**Authors:** Mauricio A Fernandez-Gonzalez

## Abstract

The high complexity of living beings represents a difficult challenge to understand how different levels of organizations are interconnected, being especially difficult to identify how certain higher levels are affected by cellular dynamics. Important advances in ecology have shown that many phenomena can be explained through a power-law relationship, being the cellular metabolism a key component in this complexity networks scaling with a regular exponent related to body size and temperature. Here, using a novel approach we estimate the energy used to synthesize the portion of the genome that codes the metabolism searching for genomic scaling rules in five different species with higher differences in body sizes. We found that the energy of this genetic portion scales in a power-law relationship related to the mass of the species analyzed.

## Introduction

The evolutionary transitions of life were characterized by astonishing increases in complexity accompanied by innovations in metabolic design. The high complexity of biological organisms has represented a huge challenge in the study of the effect of changes in one level of biological organization to other levels (Brown et al., 2004). However, advances in the ecology field place the metabolism as the key characteristic that allows using principles of physic, chemistry, and biology to link individuals with higher levels of organizations relating the metabolism with body size and temperature obeying this relationship to an exponential function that can be expressed in simple terms:

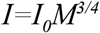

Where *I* is the metabolic rate, *M* is the body mass as the independent variable, *I*_*0*_ a normalization constant, and *b is* the scaling or allometric exponent (Kleiber, 1932). This expression establishes a power-law relationship, a ubiquitous function in physical, biological and social phenomena (Marquet et al., 2005), representing the scaling of a special case of power-law relation where the body size or temperature is the dependent variable. The relationship between body size, temperature, and metabolic rate is one of the most documented scaling relations in biology, opening a fruitful new study area that recognizes the higher complexity of biological systems and achieves to describe in simple terms his relations. The study of scaling has allowed the development of important theories on the ecology of metabolism as metabolic theory of ecology (Brown et al., 2004) and ecological stoichiometry theory (Sterner & Elser 2002).

Although scaling relationships are common at different levels of organization such as from individual to population or even ecosystems, however, works about how lower levels affect the scaling are scarce, subtracting generality to the relationships. One of the approaches that bring the phenomenon to the cellular level is the principle of stoichiometric invariance (Redfield 1958) and his emphasis on the RNA content. This principle which establishes that the stoichiometry C:N:P of biological elements yield a trade-off with the organisms and the environments, therefore the dynamics from individual to community-level, would require linking element fluxes to energetics (Redfield 1958). Due to the fractions of metabolic and structural biomass in the organisms have narrow variation in the elements that compose them, the principle of stoichiometric that can be applied to the cellular components.

Phosphorus is a key chemical element when energy and biomass are transferred between trophic levels, establishing a link between fluxes and energetics (Reiners, 1986). The variation of phosphorus concentration in organisms can be an effect variation in the concentration of P-rich ribosomal RNA (Elser et al., 1996). In higher growth rates, the RNA density is constant, because the whole-body concentration is an effect of the densities of this ribosomal RNA in a P mass fractions *c.* 0.09g P g^−1^, comprising these ribosomes an 85% of RNA. In this scenario of RNA constant concentration, the genome seems to be the unique relevant micro-level to find the first steps to the scaling, a starting point with the body size as the independent variable.

Since the genome is a static compound of the organisms in the ontogeny, therefore in the ontogeny the temperature has no effect over the genome, establishing a genomic termical invariance that does not affect the power-law function that describes the phenomenon of scaling (Lynch & Marinov, 2015). Although the cost of expression can have an effect on evolution, there are limits to this expense determined by scaling phenomena, which describe a fixed proportion of variation, transforming the organisms into a controlled system (Wagner, 2005), especially recognizing that the genetic code is nonrandom (Klump, Völker & Breslauer, 2020).

According to Lynch & Marinov (2015) each nucleotide and ribonucleotide have the same energy cost with slight variation (|~50 ATP), therefore the energy cost of synthesizing a gene depends only on the large of the genomic sequence from which the transcript is synthesized. This approach allows us to build a novel simple model to estimate the cost at the genomic level searching for a scaling relationship between genome energy related to metabolism and body-size. To analyze only the metabolism energy, we use the fraction of the genome that is translated to ribosomal compounds taking advantage of high throughput sequencing technologies to analyze five datasets of whole-proteins from five different species with largest differences in body sizes. To our knowledge, this is the first work that enlaces DNA with metabolic energy in a scaling relationship with the body-size.

## Methods

To analyze scaling in organisms with different body size, we use public data of the sequence of whole-protein from five different species: *Escherichia coli* [GenBank: U00096.3] (Denis et al., 2017), *Daphnia magna* [wfleabase: dmagset7finloc9b.puban] (Colbourne, Singan & Gilbert, 2005), *Homo sapiens* [GENCODE release 36, GRCh38.p13] (Frankish et al., 2019), *Mus musculus* [GENCODE release M25, GRCm38.p6] (Frankish et al., 2019) and *Balaena mysticetus* [Bowhead whale genome resource] (Keane et al., 2015). To build our model, we used the sequence of proteins of the five species, to count the number of codon that allows us to obtain the cost per ribonucleotide translated, hereafter the genomic theoretical base cost or *Tg*. Due to the cost of nucleotide and ribonucleotide are constant, that goes around ~ 50 ATP (Lynch & Marinov, 2015), the *Tg* per specie can be calculated using the following formula:

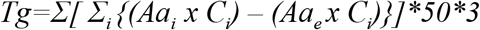

Here *Σi* is the summation of cost of synthesizing all amino acids code from gene *i* transcribed, *Aa*_*i*_ amino acids transcribed, A*ae* are the essential amino acids, *C*_*i*_ is the cost of synthesizing the *Aa*_*i*_ and A*a*_*e*_ according to Akashi and Gojobori (2002). The essential amino acids for each specie was: R-H-I-L-K-M-F-T-W-V to *E. coli* (Anderson & Jackson, 1958), R-W-H-I-L-F to daphnia, R-H-I-L-K-M-F-T-W-V to mouse (John & Bell, 1976), L-I-V-K-T-M-W-F-H to human (Reeds, 2000) and R-H-I-L-K-M-F-T-W-V to Bowhead whale (Ostrowski & Divakaran, 1989). Using this approach to estimate the genomic energy we graph the body size of the species versus the *Tg* normalizes by mass, to identify the shape of the curve to identify if the data follow a power-law relationship indicative of scaling.

Later, to estimate the fit of the data to the allometric model and once quantified the energy as *Tg*, this measure must be modeled according to allometric expression (Brown et al., 2004), which correct the metabolic rate (*I)* by the joint effects of body size (*M*), and temperature (*T*), this relation can be described as:

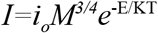

Where *M* is the body mass, *i*_*0*_ is a normalization constant independent of body size and temperature, *E* is the activation energy, *K* is Boltzmann’s constant and *T* is the absolute temperature in Kelvin. The temperature does not play a role in the energetic of the genome in the ontology, by the invariance of the code, being negligible as *E* and *K* due to and each ATP counted in *Tg*, includes in itself factors as E and K. Thus, the scaling at the genomic level, without considering a normalization constant, can be expressed as:

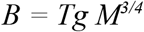

In this expression modified from Brown et al. (2004), *B* is the mass-specific *Tg* and *M* is the mass was approached using the weight in Kg of the five species: *Escherichia coli* (*1e*^−*12*^Kg) (Sajed et al., 2015)*, Daphnia magna* (*1e*^−*8*^Kg) (Simčič, & Brancelj, 1997), *Mus* musculus (*0.02 Kg*) (Cajuday & Pocsidio, 2010)*, Homo sapiens* (72 Kg) (Levine, 2006) and *Balaena mysticetus* (100000 Kg) (Keane et al., 2015). Finally we normalize the *Tg* by dividing it by the body size in kg to make the magnitudes comparable and to construct a linear model to calculate correlation and find the slope of the curve to estimate the scaling exponent using a linear model. This analysis allows us to compare the slope with the allometric exponent to estimate the model fit, finally the correlation between axis data was calculated with Pearson's product-moment correlation. All the calculations were made using R software (R Core Team, 2014).

## Results

The relationship between the body size (*M*) and the *Tg* normalized by mass for the five species follow a of power-law slope (Fig. 1) a result indicative of scaling where the organisms with less body size expend have higher *Tg* than the organisms with higher body sizes for the same quantity of mass. This result suggests that the small es species expend more energy per code to convert it into proteins.

**Figure 1.**
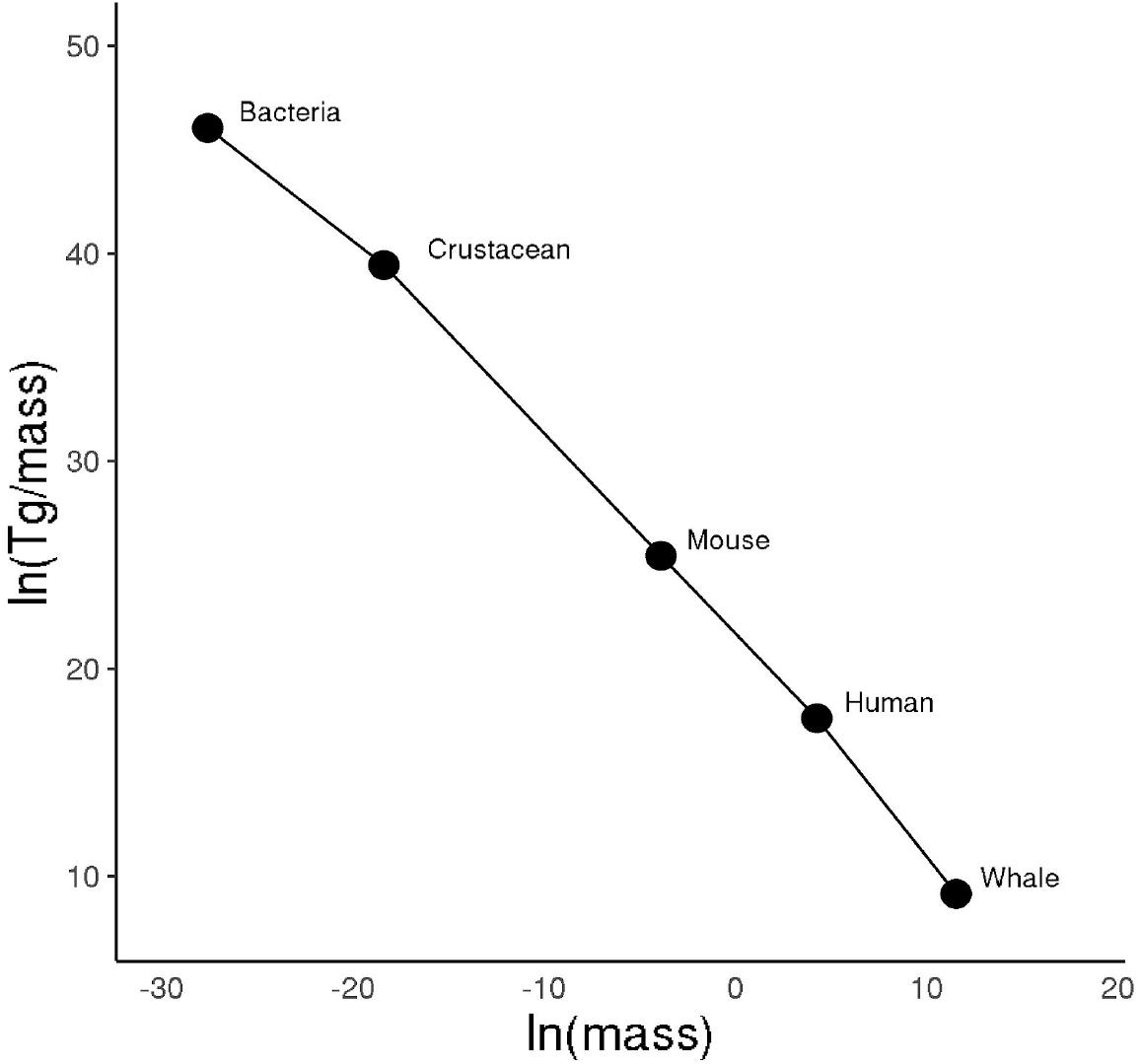
Relationship between body size and Tg normalized by mass. Energy and mass dependence at genomic level expressed as Tg for five species with huge differences in body size, both data come from five species: Escherichia coli, Daphnia magna, Homo sapiens, Mus musculus, and Balaena mysticetus. Y-axis shows Tg in the number of ATP normalized by mass in Kg. The relationship shows a power-law curve (r 2= −0.997) that is indicative of scaling. The organisms with lower body size expends higher Tg per mass unit.

The correlation between body size shows a r^2^= 0.998 and the linear model indicate a slope of 0.94 (Fig. 2), a value near to the exponent of scaling from Kleiber’s law, which indicates that the metabolic rate of organisms scales to the ¾ power of the organism mass. Both results support the idea that the scaling phenomenon would start at the level of genetic code.

**Figure 2.**
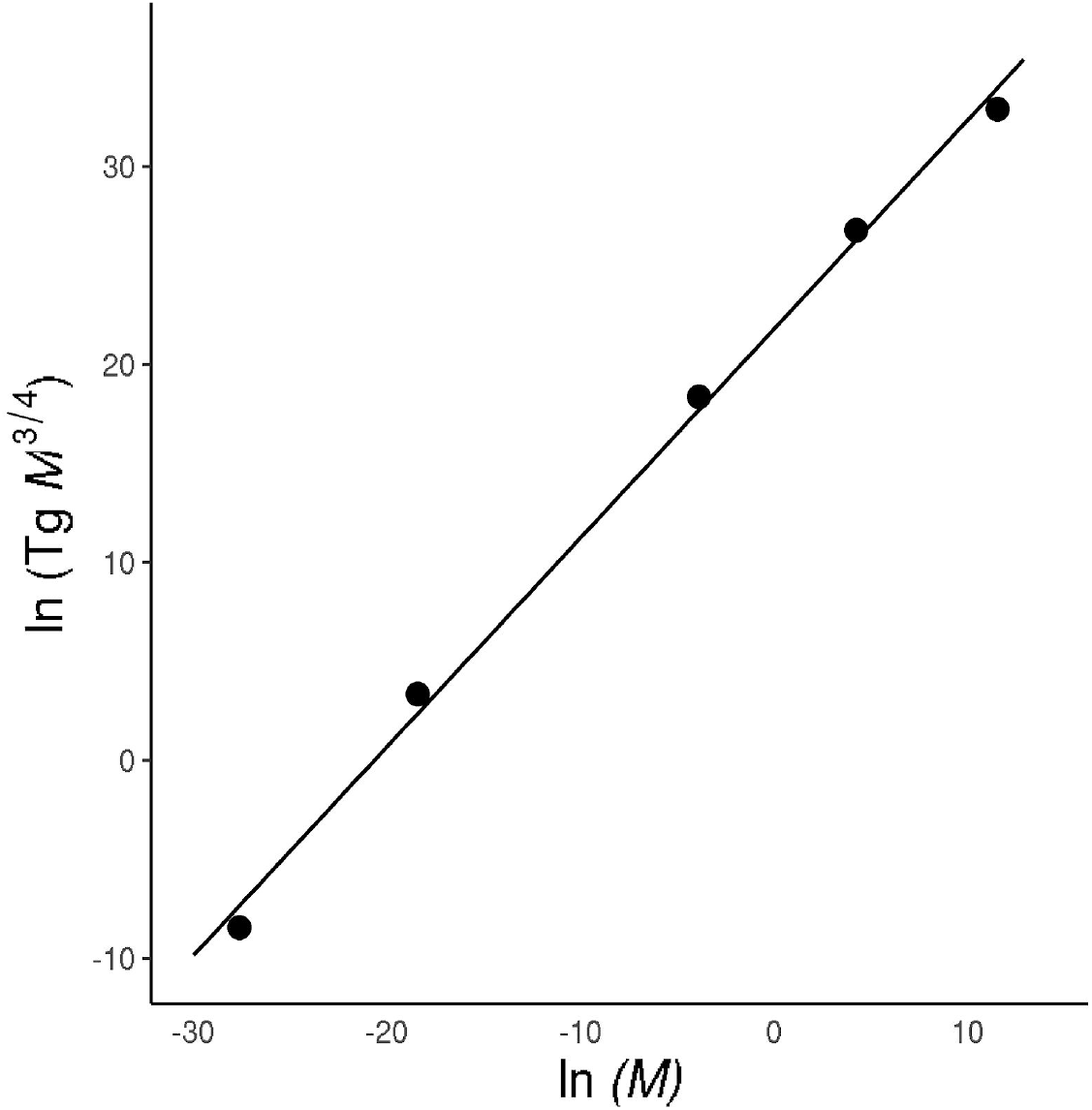
Scaling relationship between body size and *Tg* corrected by mass. Energy and mass dependence at the genomic level for five species with huge differences in body size. Y-axis shows the mass-corrected *Tg* in the number of ATP/Kg3/4. The observed slope 0.94 is close to the predicted allometric exponent of ¾, with a r^2^= 0.998. Theoretical base cost is the cost of synthesis in number ATP molecules of ribonucleotides translated to proteins and body sizes are the weight in Kilograms, both from five species.

## Discussion

Whole-genome energy of metabolism expressed as *Tg* follows power-law distribution and this energy decreases with the increase in body size across the five species analyzed (Fig. 1-2). This scaling relationship shows that the organisms with lower body size expend higher genomic energy in *Tg* form per mass unit. The differences between the slope calculated and the allometric exponent can be an effect of the precision of the sequence techniques to obtain high throughput sequencing data, the quantity of data analyzed and quality of annotation of the sequences. However, these differences can be negligible considering the r^2^ = 0.997 calculated, possibly more precision on techniques of data collection they would only add slight differences bringing the slope even closer to an allometric exponent, without losing sight that metabolic rate can be influenced by different constraints in different life forms (DeLong et al., 2010).

It is easy to fall into the discussion of the effect of a nucleotide more or less in the code can generate big changes, however, it is necessary to take into account that any energy analysis methodology like ours, must take into account that the energy cost of genomes with more and less complex structures are similar because always follow a power-law relationship (Lynch and Marinov, 2015), where the growth requirements per cell scale with cell volume in a power-law relationship with highly significant regression (r^2^ near to 1) indicating a linear relationship between the energetic requirement for growth and cell volume that involve for example, the same pattern when compare prokaryotic and eukaryotic cells (Lynch and Marinov, 2015, Chiyomaru & Takemoto, 2020). This example, implies that any overweight of importance of specific components of genomes in the energy expenditure invokes an, unlike variable metabolic scaling. For example, if a simple mutation that incurs an extra energetic cost, may be compensated for by other adaptive effects (Ilker & Hinczewski, 2019), and the phenomenon returns to scale into a power-law relationship. Therefore the development of an energetic genetic model that uses global measures as RNASeq with more or less specificity does not affect the result of the scaling phenomenon observed.

The negligible importance of RNA concentration in scaling and the highly importance of genomic to explain the higher levels of organizations, can be related to the use of codons in organisms that connect replication and transcription to translation, allow to evolve more rapidly toward a thermodynamically driven code (Klump, Völker & Breslauer, 2020). Darwin establish that (Darwin, 1859) if a phenotypic characteristic gives a survival advantage this characteristic persists through generations, the ubiquity scaling phenomenon is not the exception as a genotypic signature that give advantageous phenotypical characteristics relates to code and not just exclusively in terms of the functionality, due to the codon usage already exist in different species, a selected code that allows the optimization of energy in thermodynamic stability (Klump, Völker & Breslauer, 2020). The constance of scaling in lower levels can be an effect of the less sensitivity of coding sequences to environmental conditions, allowing them to keep their function stable (Klump, Völker & Breslauer, 2020) in a necessary stability and advantageous system.

Previous works have proposed a power-law at the genomic level, but in these approaches the code of DNA is analyzed as natural language, concluding that coding and noncoding regions follow a power law (Luscombe et al., 2002). These behaviors are also found in sub-regions of the genome, as pseudogene and pseudomotif, intergenic regions and the number of expressed mRNA transcript (Luscombe et al., 2002), confirming the ubiquity of scaling genomic approaches. Due to the cost of nucleotide and ribonucleotide are constant (Lynch & Marinov, 2015) and the invariance of the genetic code in the ontology of organisms, the scaling-genomic phenomenon found here also would apply undoubtedly to direct analysis of DNA sequences.

To our knowledge, this is the first work that link genomic code with energetic count searching for genetic-metabolic scaling relationship with the body-size, taking advantage of the benefits of using data from high throughput platforms, that allow us to get closer to the complete genome, transcriptome, proteome and metabolome of the organisms. Even though the genome is almost invariable, and to be addressed in the future, it would be relevant to include more high throughput data from new species with different body sizes, in a standardized pipeline that also includes replicates to these data, to get a more accurate slope to the linear model.

